# ECS1 and ECS2 regulate polyspermy and suppress the formation of haploid plants by promoting double fertilization

**DOI:** 10.1101/2022.01.20.476184

**Authors:** Yanbo Mao, Thomas Nakel, Isil Erbasol Serbes, Dawit G. Tekleyohans, Saurabh Joshi, Thomas Baum, Rita Groß-Hardt

**Affiliations:** University of Bremen, Centre for Biomolecular Interactions, 28359 Bremen, Germany

## Abstract

In flowering plants, the number of pollen tubes that provide sperm cells to the female gametes is restricted by a pollen tube block. This safeguard mechanism is only activated after successful fertilization of both female gametes and involves the disintegration of pollen tube attracting synergid cells. Yu and coworkers previously reported that the endopeptidase ECS1 and ECS2, which are secreted by fertilized egg cells, prevent the attraction of supernumerary pollen tubes by cleaving the pollen tube attractant LURE1^1^. Here we report on an earlier defect in *ecs1 ecs2* mutants that is manifested by single rather than double fertilization of either egg or central cell. The defect is accompanied by a delay in synergid disintegration providing an alternative explanation for the extra pollen tubes observed in the double mutant. These results are corroborated by our finding that *ecs1 ecs2* plants segregate both, haploid as well as polyspermy derived triploid plants.

---

Yu et al.^1^ investigated two aspartic endopeptidases, ECS1 and ECS2, that are secreted from the egg cell towards the filiform apparatus upon sperm-egg fusion. In addition, they provide a compelling set of evidence demonstrating that both ECS1 and ECS2 can cleave specific variants of LURE1. The authors observed normal rates of fertilization of both female gametes in *ecs1 ecs2* double mutants and they concluded that it is the LURE1 cleaving function of *ECS1* and *ECS2* that accounts for 10 % supernumerary pollen tubes attracted by *ecs1 ecs2* ovules. In contrast to the work by Yu et al., under our growth conditions *ecs1 ecs2* double mutants do exhibit a fertilization defect, which is reflected by 13 % abnormal seeds (Fig. 1a, Extended Data Fig. 1a). In the following we will provide data that suggest single fertilization as an alternative course for the observed supernumerary pollen tube attraction in *ecs1 ecs2* and we show that lack of ECS1/ECS2 function can induce haploidy as well as polyspermy derived triploids.

**Fig. 1.**
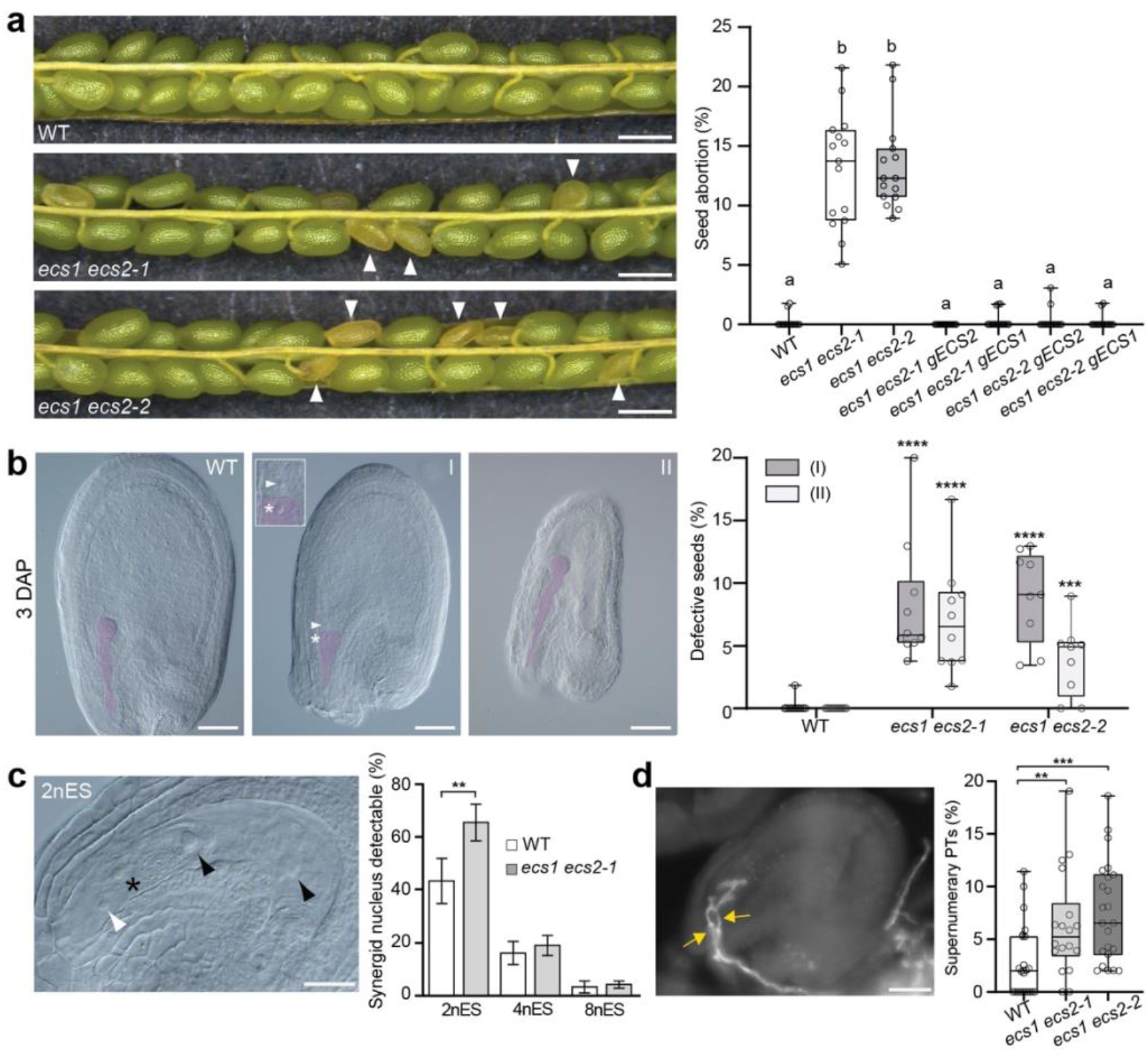
ECS1 and ECS2 promote double fertilization and prevent the formation of haploid plants. **a**, Silique of wild type and *ecs1 ecs2* mutants and quantification of seed abortion in different genotypes (n > 800 for each genotype). Circles represent data recovered from individual siliques. Arrowheads, aborted seeds. Different letters show significant difference, p < 0.0001, *F* =106.0, by one way ANOVA with a Tukey multiple comparison test. **b**, Cleared whole mount and quantitative analysis of *ecs1 ecs2* mutant seed categories 3 days after pollination (DAP): In comparison to wild-type like seeds containing embryo and endosperm, *ecs1 ecs2* mutants segregate seeds without embryo (I) and seeds containing no or retarded endosperm (II). (n=655/540/522 for wild type/ *ecs1 ecs2-1/ ecs1 ecs2-2* respectively). Circles represent data recovered from individual pistils. Embryo and unfertilized egg cells were false coloured in purple. Asterisk, unfertilized egg nucleus; arrowhead, endosperm nucleus. Similar results for **a** and **b** were obtained in independent experiments by a different scientist (data shown in Extended Data Fig. 1a and d). **c**, Frequency of seeds containing a synergid nucleus at different endosperm stages (n for 2nES= 67/72, 4nES=143/158 and 8nES= 179/141 for wild type/ *ecs1 ecs2-1* respectively). Data were generated by two different scientists and combined. The image shows the second synergid nucleus at two-nucleate endosperm stage. Endosperm nuclei, black arrowhead; synergid (syn) nucleus, white arrowhead; zygote, asterisk. **d**, Occurrence of polytubey in *ecs1 ecs2* mutants 20 hours after pollination (HAP) (n=1039/835/1070 for wild type/ *ecs1 ecs2-1/ ecs1 ecs2-2* respectively). Circles represent data recovered from individual pistils. The image indicates two pollen tubes (yellow arrow) targeting an embryo sac. Data in **a**, **b**, **d** are represented in box and whisker plot, center line, median; bottom and up, 25^th^ and 75^th^ percentiles. Whisker, the minimum and maximum. Data in **c** indicate mean ± SD. Two-tailed Mann Whitney comparison test between wild type and mutants, ** p < 0.01; *** p < 0.001; **** p < 0.0001. Scale bars, 500 μm (**a**), 50 μm (**b**) and (**d**), 20 μm (**c**).

To investigate whether the reproductive defect was causally linked to ECS loci, we first generated a second *ecs1 ecs2* double mutant using an independent *ecs2* allele, which exhibited similar seed development defects (Fig. 1a, Extended Fig. 1a). Second, we introduced *pECS1:gECS1-YFP* and *pECS2:gECS2-YFP* into the *ecs1 ecs2* double mutants and demonstrated that each of the constructs restored fertility (Fig. 1a, Extended Fig. 1a). Our results thus indicate that the seed defect in *ecs1 ecs2* double mutants is caused by loss of ECS1 and ECS2. To determine whether the defect was due to a role of ECS1 and ECS2 in the male or female reproductive tissue, we conducted reciprocal crosses between the double mutant and wild type. While the introduction of the mutant alleles from the female crossing partner recapitulated the defect, the reciprocal cross yielded fertile seeds, suggesting that the defect was of a female origin (Extended Data Fig. 1b). To further narrow down the defect, we performed a comprehensive analysis of cleared whole mounts of mutants and wild type at different developmental stages. Prior to fertilization, *ecs1 ecs2* female gametophytes exhibited a morphology similar to that of wild type (Extended Data Fig. 1c). But, three days after pollination, we observed 8.6 % of seeds that failed to develop an embryo and 5.5 % of seeds that exhibited defects in endosperm development (Fig. 1b, Extended Data Fig. 1d). These observations were indicative of single fertilization in either of the two female germ cells. Previous work has uncovered that defective or incomplete gamete fusion induced, for example, by an insufficient number of fertilization-competent sperm^2–8^, triggers a fertilization recovery program. The mechanism includes transient suspension of second synergid disintegration and concomitant attraction of supernumerary pollen tubes^9–13^, which we accordingly expected to see in the mutant.

When inspecting young *ecs1 ecs2* seeds at the two-nucleate endosperm stage, we indeed found a significant increase of synergid-containing seeds and supernumerary pollen tube attraction as compared to wild type (Fig. 1c, d). Overall progression of seed development between mutant and wild type was comparable and *ecs1 ecs2* synergid nuclei eventually degenerated (Fig. 1c, Extended Data Fig. 1e). It is noteworthy that the frequency of single fertilization can account for the polytubey observed by Yu et al. and in this study. In addition, our observations suggest that this female defect in gamete fusion cannot efficiently be rescued by the fertilization recovery mechanism.

Last but not least we asked whether the reproductive defect was reflected in the offspring of *ecs1 ecs2* double mutant. Upon inspection of 200 *ecs1 ecs2* double mutant offspring, we identified 7 plants with short siliques and small flowers. Using flow cytometry, we found that all 7 plants contained a haploid genome (Fig. 2a, b, Extended Data Fig. 2), suggesting parthenogenetic activation of an unfertilized egg cell. Several factors regulating gamete fusion have been described^2–7,14^. Among them, only the sperm specific AtDMP8 and AtDMP9 have been implicated in haploid induction^15^. It will be an interesting task for the future to determine whether other gamete fusion defective mutants similarly give rise to haploids or whether this phenotype uncovers an additional function of ECS1 and ECS2. Second, we asked whether conversely also polyspermy, the fusion of an egg by more than one sperm was enhanced. We have previously established a polyspermy detection assay (HIPOD). The assay consists of two pollen donors, which contain the individual elements of the two-component system, *mGAL4-VP16/pUAS*^16^. The elements control expression of an herbicide resistance conferring *BAR* gene tagged to a yellow fluorescent protein (*BAR-YFP*) such that resistance is only established if both paternal constructs are combined in a single egg cell^16^. We pollinated both, *ecs1 ecs2-1* double mutants and wild type with pollen from both fathers. For wild type we recovered 24 triparental plants out of 96,590, corresponding to a polyspermy frequency of 0.05 %. By comparison, we detected 53 triparental plants out of 77,441 in *ecs1 ecs2-1* double mutants, corresponding to a polyspermy frequency of 0.15 % (Fig. 2c-f). We confirmed that this phenomenon was indeed due to ECS1 and ECS2 by introducing a *pECS1∷gECS1-YFP* rescue construct, which restored triparental frequencies to wild type levels (Fig. 2f). ECS1 and ECS2 are hence the first molecular factors demonstrated to affect polyspermy in plants.

**Fig. 2.**
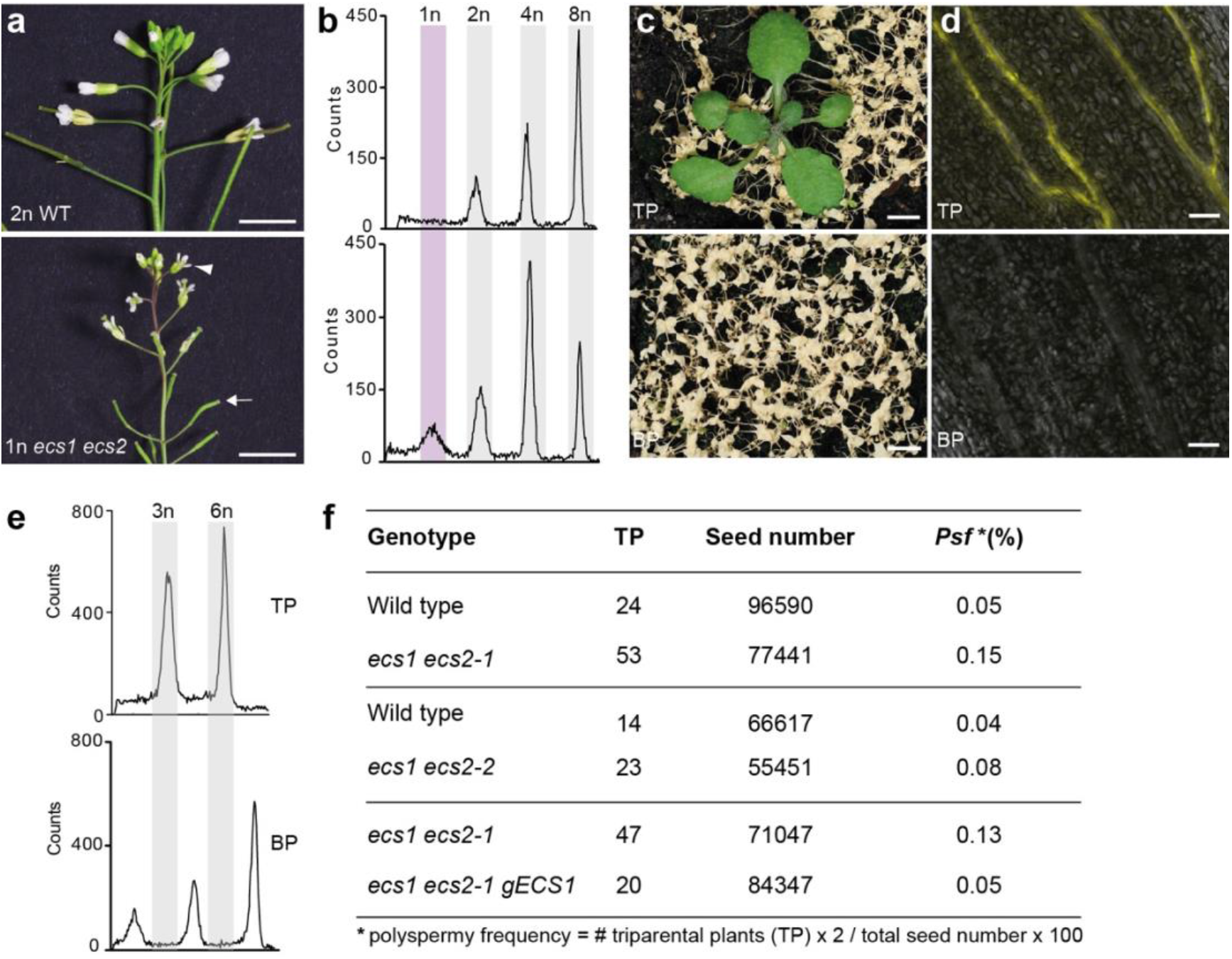
*ecs1 ecs2* double mutants segregate haploid and triparental offspring. **a** - **b**, Inflorescence **(a)** and flow cytometry **(b)** of diploid (2n) wild type and haploid (1n) *ecs1 ecs2* plant (n=200, see also Extended Data Fig. 2). Brightness was manually enhanced with Adobe Photoshop in **(a)**. The arrowhead and arrow point at a small flower and an undeveloped silique, respectively. **c - e**, Herbicide treatment (**c**), YFP fluorescence (**d**), and ploidy analysis (**e**) of triparental triploid plants (TP), and biparental diploid plants (BP). (**f**) Polyspermy frequency in *ecs1 ecs2* mutants compared to wild type and the complementary transgenic line *ecs1 ecs2 pECS1∷ gECS1-YFP*. Scale bars, 5 mm (**a**) and (**c**), 50 μm (**d**).

Taken together, we have shown that ECS1 and ECS2 regulate offspring genome size via haploid and polyspermy induction. In addition, our observation of evident single fertilization and a temporal delay of second synergid disintegration provides an alternative explanation for the attraction of supernumerary pollen tubes observed both by Yu et al^1^. and in this study.

## Acknowledgments

We thank members of the Groß-Hardt laboratory for valuable comments on the manuscript. This work is supported by the European Research Council (ERC Consolidator Grant “bi-BLOCK” ID. 646644 to R.G.).

## Author contributions

Y.M., T.N., I.E.S, D.T., S.J., T.B., and R.G. designed the experiments. Y.M., T.N., I.E.S, D.T., S.J., and T.B. performed and analysed the data. All authors discussed the results and contributed to the manuscript.

## Material and Methods

### Plant material and growth conditions

All experiments were performed with *Arabidopsis thaliana* L., ecotype Col-0. The seeds were sown on soil. 2 days after stratification at 4 °C, they were transferred and germinated in a Conviron MTPS growth chamber under long-day conditions (16 hr light/8 hr dark) at 23 °C. Plants were transferred into 18 °C after bolting. T-DNA insertion mutants SALK_021086 (*ecs1*), SALK_090795 (*ecs2-1*), SALK_036333 (*ecs2-2*) were obtained from European Arabidopsis Stock Center (NASC) (Nottingham, UK).

### Plasmid construction and plant transformation

The promoter and gene fragments of *ECS1* and *ECS2* were amplified from Arabidopsis Col-0 by using YM76s/as and YM78s/as and introduced into *pUAS∷BASTA-YFP*^16^ using AscI/NotI restriction sites. The resulting constructs were then transformed into *ecs1 ecs2* double mutants by floral dip^17^. Primer sequences used for genotyping and plasmid construction are listed in Supplementary Table 1.

### Histology and microscopy

For the analysis of mature female gametophytes, the oldest closed flower buds were emasculated and harvested two days later. For the analysis of seed development, the emasculated flowers were manually pollinated and collected as specified in the figure legends. For whole mount clearings, samples were treated with acetic acid/ethanol (v/v = 1:9) overnight, washed in a 80%, 70% and 50% ethanol series and mounted in chloral hydrate:glycerol:water (8:2:1; w:v:v)^18^. For each experiment, more than 5 emasculated flowers or pollinated pistils were analysed. With regard to the determination of persistent synergid cell, more than 10 pistils 19-21 hours after pollination (HAP) were scored and the data from the pistils containing less than 10 fertilized ovules were excluded. Cleared whole mounts of ovules and seeds were analysed under Zeiss Axioscope (Zeiss, Oberkochen, Germany) and the images were captured by Canon PowerShot G10 camera. Mature siliques were dissected and observed under Leica S6E or S8apo stereomicroscope (Leica, Germany).

For aniline blue staining, pistils were collected 20 HAP and processed as described previously^2^. This experiment was analyzed for each genotype with two technical replicates. Supernumerary pollen tube attraction was examined using an epifluorescence inverted microscope (Leica DMI6000b) with DAPI filter.

### Triparental plant screening

To determine the occurrence of polyspermy-induced triparental plants, HIPOD screening was applied as previously described^16^. 2-3 closed flower buds per inflorescence were emasculated. Two to three days after emasculation pollen grains collected from plants of *pRPS5a∷mGAL4-VP16/+* and *pUAS∷BAR-YFP /+* were applied onto the stigmatic surface. The resulting mature seeds were sown on soil and the plants were subjected to herbicide treatment. The triparental status of herbicide-resistant plants was confirmed by flow cytometry, fluorescence microscopy inspection and PCR targeting of the *GAL4* and *UAS* loci.

### Ploidy analysis

Leaves of *ecs1 ecs2* plants or herbicide-resistant plants from HIPOD screenings were subjected to flow cytometry as previously described^16^.

### Statistical analysis

Data sets were processed in Microsoft Excel. Bar charts, box and whisker plots as well as all statistical tests were generated using Microsoft Excel or GraphPad Prism.

---

**Extended Data Fig. 1.**
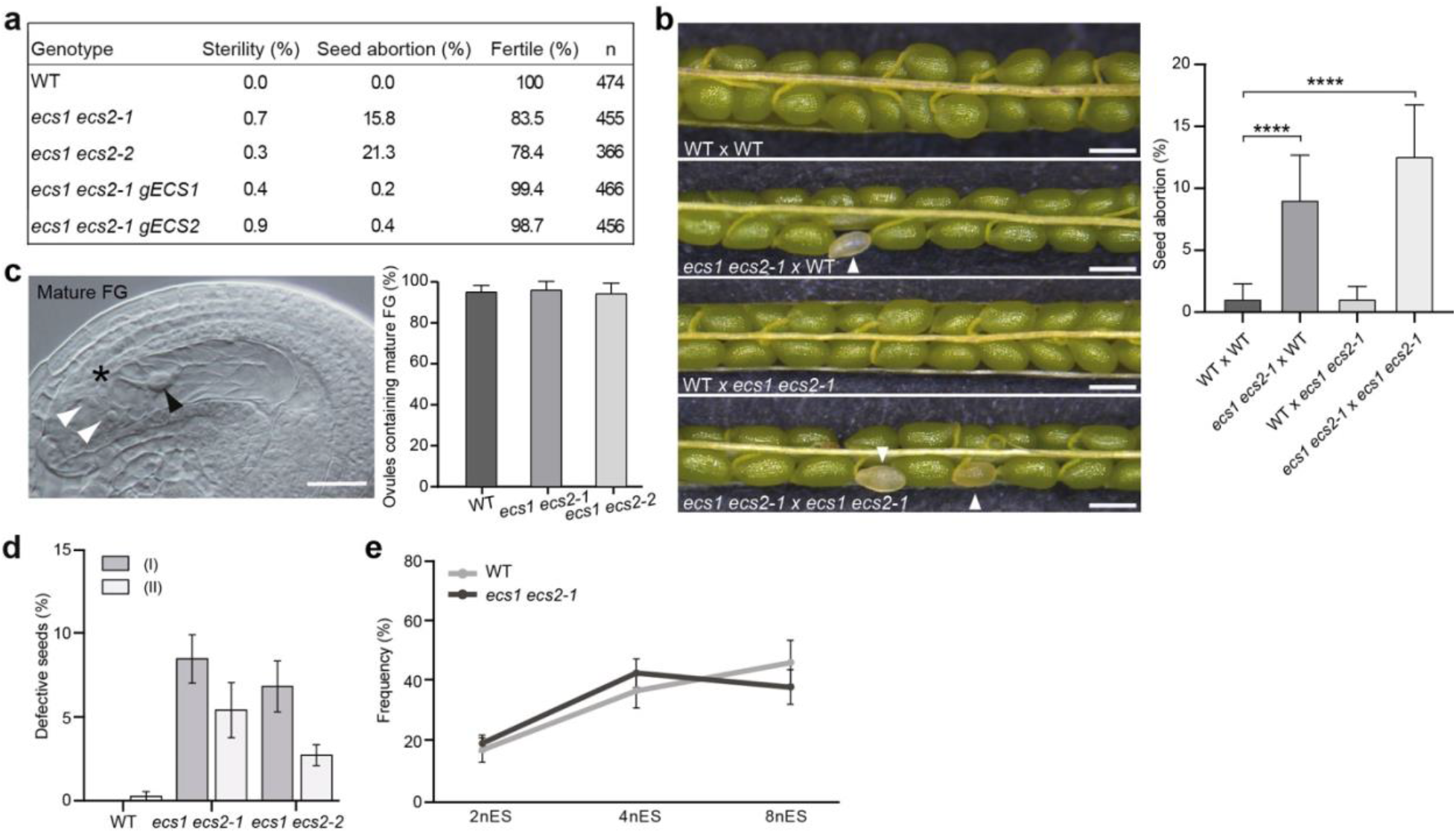
The seed defect in *ecs1 ecs2* is of a female origin. **a**, Frequency of seed abortion in different genotypes. **b**, Abnormal seed formation in plants resulting from crosses between different genotypes (n= 614/375/382/395 for WT × WT/*ecs1 ecs2-1*×WT/ WT × *ecs1 ecs2-1*/*ecs1 ecs2-1*× *ecs1 ecs2-1*, respectively). Arrowheads, aborted seeds. Similar results were obtained by an additional scientist (data not shown). Data are mean values ± SD; **** p < 0.0001 (Mann Whitney comparison test). **c**, Frequencies of ovules containing mature FGs (n=419/398/424 for WT/ *ecs1 ecs2-1*/ ecs*1 ecs2-2*, respectively). Central cell nucleus, black arrowhead; synergid (syn) nucleus, white arrowhead; egg nucleus, asterisk. **d**, Quantitative analysis of *ecs1 ecs2* mutant seed categories 3 DAP: In comparison to wild-type like seeds, seeds without embryo (I) and seeds containing no or retarded endosperm (II) were observed (n=356/295/366 for wild type/ *ecs1 ecs2-1/ ecs1 ecs2-2* respectively). **e**, Developmental stage of WT and *ecs1 ecs2-1* seeds 20 HAP: Categories detected are two-, four- and eight-nucleate endosperm staged seeds (2nES, 4nES and 8nES) (n=389/371 for WT/ *ecs1 ecs2-1*, respectively). No statistically significant differences detected on the basis of either student’s t-test or Mann Whitney comparison test.

**Extended data Fig. 2.**
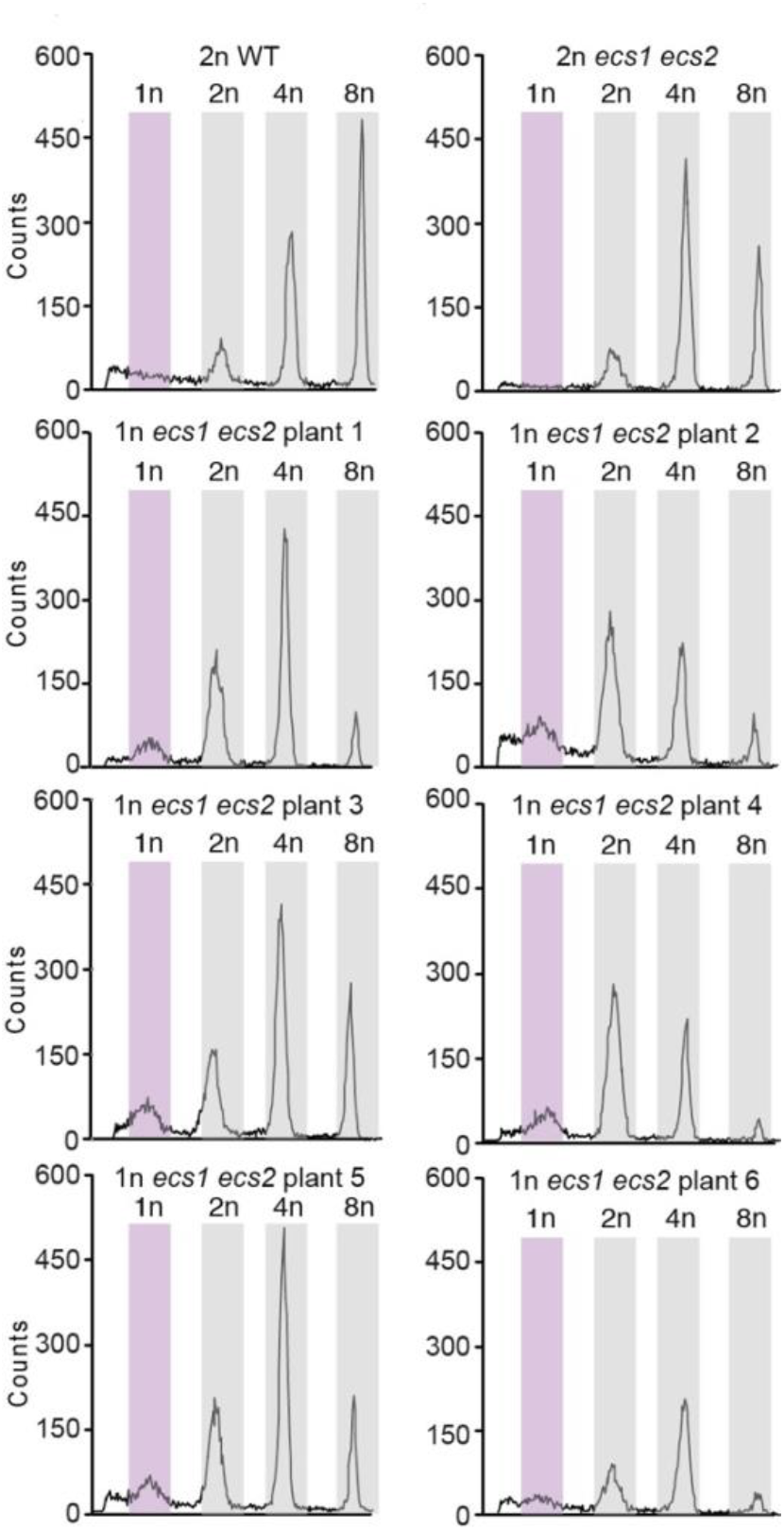
Ploidy analysis of phenotypically conspicuous *ecs1 ecs2* double mutant offspring. Flow cytometry analysis of diploid wild-type, diploid (2n) *ecs1 ecs2*, and six haploid (1n) *ecs1 ecs2* plants. The haploid profile is highlighted by purple color.

