## Supplementary Table 1 for "ECS1 and ECS2 regulate polyspermy and suppress the formation of haploid plants by promoting double fertilization"

**Primer sequences used for genotyping and plasmid construction**

| Oligos | Sequence (5'-3') | Application |
|---|---|---|
| YM66s<br>YM66as | TTCACAACCAAAACCGATCTC<br>AAAACGCTAGCTCTTAACGGC | Genotyping of SALK_021086 (*ecs1*) |
| YM67s<br>YM67as | TCGAAGGTACCACCGTTATTG<br>CGGCCAAAAGTACTCTCAGTG | Genotyping of SALK_090795 (*ecs2-1*) |
| YM68s<br>YM68as | CAGTCGTTCAGCAAAGGAGAC<br>ATTGGTAGATTGGTGACACGC | Genotyping of SALK_036333 (*ecs2-2*) |
| YM76s (AscI)<br>YM76as (NotI) | GGCGCGCCGAGATTTGGGAAATGTGCAAT<br>GCGGCCGCTAAGTTCCCGGAGCAATCCAT | Amplification of genomic region of *ECS1* |
| YM78s (AscI)<br>YM78as (NotI) | GGCGCGCCCCATCCTCATGACAGTACCTGGA<br>GCGGCCGCCAAATTCGCAGAGCAATCCAT | Amplification of genomic region of *ECS2* |
| TN12s<br>TN12as | ATTTAATTAAATGAAGCTCCTGTCCTCCATCGA<br>ATGCGGCCGCCTACCCACCGTACTCGTCAATTC | *GAL4* detection |
| TN26s<br>TN26as | GCAGGAGTGGACGGACGACCTCGTCCGTCTG<br>TCACGAACTCCAGCAGGACCATGTGATCGCG | *UAS* detection |
